# Bacterial sequences detected in 99 out of 99 serum samples from Ebola patients

**DOI:** 10.1101/039107

**Authors:** Marina Manrique, Eduardo Pareja-Tobes, Eduardo Pareja, Pablo Pareja-Tobes, Raquel Tobes

**Author notes:** Era7 Bioinformatics, Plaza Campo Verde 3, Atico. Granada 18001. Spain. Corresponding author: Raquel Tobes.

## Abstract

Evolution and clinical manifestations of Ebola virus (EBOV) infection overlap with the pathologic processes that occur in sepsis^1^. Some viruses certainly compromise the immune system, leading to a breach in the integrity of the mucosal epithelial barrier, thus allowing bacterial translocation^2, 3^. Guided by these facts, we wondered if bacteria could be involved in the pathogenesis of some of the septic shock-like symptoms typical of EBOV infected patients, something that could have a dramatic impact on the design of new treatment approaches. We decided to search for bacteria in available EBOV patient sequence datasets. Given that EBOV is an RNA virus and that, hence, some NGS sequencing experiments carried out to sequence the EBOV genomes were RNA-Seq experiments, we thought that, if there were any bacteria in patient serum, at least some bacterial RNA might probably be detected in the sequenced material from Ebola patients. Thus, we searched for bacteria in a RNA-Seq public dataset from 99 Ebola samples from the last outbreak^4^, and surprisingly, in spite of the certainly suboptimal experimental conditions for bacterial RNA sequencing, we found bacteria in all of the 99 samples.

## RNA-Seq dataset from Ebola patients

The datasets with the illumina reads of the samples from the Ebola outbreak published^4^ were retrieved from the SRA. For each sample listed in the Supplementary Table S2 of the article^4^ the Experiment sample (SRS) information and links to its SRA experiments (SRX) and runs (SRR) were retrieved using the NCBI Batch Entrez platform (http://www.ncbi.nlm.nih.gov/sites/batchentrez).

We analyzed a total of **166.710.398** reads from this 99 RNA-Seq dataset of the sequences obtained from serum from Ebola patients. Table 1 shows the number of reads analyzed and the reads assigned to bacteria in each sample. SRA IDs corresponding to the 99 Datasets^4^, description of Bioinformatics methods and additional data about statistics and frequency distribution of detected bacteria are available as Supplementary Material. We searched for the presence of bacterial 16S rRNA sequences. The reads for each sample were used as query sequences for BLASTn against a custom 16S database using MG7 metagenomics profiling tool^5^. The database was built with the GenBank sequences corresponding to the sequences published in the RDP database (https://rdp.cme.msu.edu/index.jsp). Based on the BLASTn results we performed a subsequent taxonomic assignment to the NCBI taxonomy using 2 different assignment paradigms: weighted Lowest Common Ancestor (LCA) and Best BLAST Hit (BBH). This analysis was done using MG7 metagenomics profiling tool^5^; this cloud-based bacterial community profiling tool (open source code available at GitHub https://github.com/ohnosequences/mg7) maps the input sequences against a database of bacterial 16S sequences using BLAST, and uses these hits to identify their taxonomic node through a BBH or a weighted LCA paradigm.

**Table 1.** Statistics of bacterial assigned reads.

| Sample ID | Analyzed reads (after filtering) | No Hit (against 16S bacterial RNA DB) | Not assigned | bacterial assigned reads (using a 16S bacterial RNA database) | % bacterial reads | outcome |
| --- | --- | --- | --- | --- | --- | --- |
| EM104 | 927027 | 912988 | 6610 | 7429 | 0.80 | Died |
| EM110 | 4892870 | 4891785 | 459 | 543 | 0.01 | Died |
| EM111 | 1464419 | 1450936 | 6025 | 7458 | 0.51 | Died |
| EM112 | 3648071 | 3644895 | 1335 | 1841 | 0.05 | Died |
| EM113 | 964090 | 949001 | 8350 | 6739 | 0.70 | Died |
| EM115 | 1222962 | 1206106 | 7259 | 9597 | 0.78 | Died |
| EM119 | 2279332 | 2270073 | 4251 | 5008 | 0.22 | Died |
| EM120 | 2160593 | 2145698 | 7130 | 7765 | 0.36 | Died |
| EM121 | 3069339 | 3065083 | 2204 | 2052 | 0.07 | Died |
| EM124-1 | 2120862 | 2113021 | 4209 | 3632 | 0.17 | Died |
| EM124-2 | 755247 | 738574 | 5950 | 10723 | 1.42 | Died |
| EM124-3 | 768610 | 762537 | 3194 | 2879 | 0.37 | Died |
| EM124-4 | 222370 | 220913 | 591 | 866 | 0.39 | Died |
| G3676-1 | 2161216 | 2151979 | 3755 | 5482 | 0.25 | Died |
| G3676-2 | 2190925 | 2186248 | 1280 | 3397 | 0.16 | Died |
| G3677-1 | 1577917 | 1577083 | 261 | 573 | 0.04 | Died |
| G3677-2 | 1834471 | 1833450 | 217 | 804 | 0.04 | Died |
| G3707 | 1545610 | 1513805 | 11116 | 20689 | 1.34 | Died |
| G3713-2 | 3256482 | 3253707 | 826 | 1949 | 0.06 | Died |
| G3713-3 | 2760034 | 2751089 | 3495 | 5450 | 0.20 | Died |
| G3713-4 | 2936648 | 2932385 | 1588 | 2675 | 0.09 | Died |
| G3724 | 3401326 | 3401088 | 116 | 122 | 0.003 | Died |
| G3735-1 | 3652776 | 3650681 | 512 | 1583 | 0.04 | Died |
| G3735-2 | 3900255 | 3899104 | 314 | 837 | 0.02 | Died |
| G3752 | 755678 | 754752 | 255 | 671 | 0.09 | Died |
| G3764 | 3566180 | 3565650 | 160 | 370 | 0.01 | Died |
| G3770-1 | 2527295 | 2517367 | 2769 | 7159 | 0.28 | Died |
| G3770-2 | 5208479 | 5199893 | 3032 | 5554 | 0.11 | Died |
| G3787 | 681519 | 667834 | 5420 | 8265 | 1.21 | Died |
| G3795 | 1099027 | 1089879 | 4185 | 4963 | 0.45 | Died |
| G3798 | 821307 | 810289 | 5054 | 5964 | 0.73 | Died |
| G3800 | 1974089 | 1946261 | 11317 | 16511 | 0.84 | Died |
| G3807 | 1599122 | 1569590 | 13024 | 16508 | 1.03 | Died |
| G3808 | 2060388 | 1979793 | 31807 | 48788 | 2.37 | Died |
| G3814 | 835800 | 812366 | 10074 | 13360 | 1.60 | Died |
| G3816 | 809251 | 770952 | 16939 | 21360 | 2.64 | Died |
| G3818 | 2044579 | 2023665 | 7953 | 12961 | 0.63 | Died |
| G3820 | 1897113 | 1876463 | 7434 | 13216 | 0.70 | Died |
| G3822 | 2542423 | 2535011 | 2886 | 4526 | 0.18 | Died |
| G3823 | 3418212 | 3409133 | 3089 | 5990 | 0.18 | Died |
| G3825-1 | 2525303 | 2495191 | 11298 | 18814 | 0.75 | Died |
| G3825-2 | 2760725 | 2755436 | 2042 | 3247 | 0.12 | Died |
| G3826 | 946841 | 934807 | 4982 | 7052 | 0.74 | Died |
| G3827 | 786789 | 776089 | 4211 | 6489 | 0.82 | Died |
| G3829 | 1434654 | 1397643 | 12567 | 24444 | 1.70 | Died |
| G3831 | 2101611 | 2091908 | 3266 | 6437 | 0.31 | Died |
| G3834 | 2015904 | 1994714 | 11613 | 9577 | 0.48 | Died |
| G3838 | 1564305 | 1556398 | 3078 | 4829 | 0.31 | Died |
| G3840 | 2382513 | 2377172 | 2774 | 2567 | 0.11 | Died |
| G3845 | 2712208 | 2705089 | 2928 | 4191 | 0.15 | Died |
| G3846 | 2307278 | 2301597 | 2204 | 3477 | 0.15 | Died |
| G3848 | 2489357 | 2484112 | 1949 | 3296 | 0.13 | Died |
| G3851 | 1917446 | 1897411 | 10276 | 9759 | 0.51 | Died |
| G3856-1 | 2189510 | 2188838 | 231 | 441 | 0.02 | Died |
| G3856-3 | 1176332 | 1159797 | 9682 | 6853 | 0.58 | Died |
| EM106 | 785650 | 774050 | 6352 | 5248 | 0.67 | Discharged |
| G3670-1 | 1603798 | 1597937 | 2049 | 3812 | 0.24 | Discharged |
| G3765-2 | 485083 | 478798 | 2320 | 3965 | 0.82 | Discharged |
| G3769-1 | 1219532 | 1208037 | 4914 | 6581 | 0.54 | Discharged |
| G3769-2 | 806309 | 798797 | 2696 | 4816 | 0.60 | Discharged |
| G3769-3 | 1262784 | 1238388 | 11790 | 12606 | 1.00 | Discharged |
| G3769-4 | 257875 | 257578 | 206 | 91 | 0.04 | Discharged |
| G3789-1 | 582942 | 570205 | 5300 | 7437 | 1.28 | Discharged |
| G3796 | 2093457 | 2081103 | 4615 | 7739 | 0.37 | Discharged |
| G3799 | 1415467 | 1386254 | 10825 | 18388 | 1.30 | Discharged |
| G3805-1 | 1231640 | 1193536 | 18366 | 19738 | 1.60 | Discharged |
| G3805-2 | 293178 | 289975 | 1334 | 1869 | 0.64 | Discharged |
| G3809 | 695337 | 672263 | 10351 | 12723 | 1.83 | Discharged |
| G3810-1 | 1007240 | 979654 | 12563 | 15023 | 1.49 | Discharged |
| G3810-2 | 769238 | 763558 | 2707 | 2973 | 0.39 | Discharged |
| G3817 | 1094044 | 1078801 | 6364 | 8879 | 0.81 | Discharged |
| G3819 | 764579 | 748395 | 6304 | 9880 | 1.29 | Discharged |
| G3821 | 1128371 | 1103076 | 10264 | 15031 | 1.33 | Discharged |
| G3850 | 480504 | 447848 | 14395 | 18261 | 3.80 | Discharged |
| G3857 | 507320 | 499773 | 3090 | 4457 | 0.88 | Discharged |
| NM042-1 | 755816 | 747548 | 3241 | 5027 | 0.67 | Discharged |
| NM042-2 | 1257035 | 1226132 | 14582 | 16321 | 1.30 | Discharged |
| NM042-3 | 439674 | 435580 | 1310 | 2784 | 0.63 | Discharged |
| EM095 | 3731730 | 3626319 | 36020 | 69391 | 1.86 | N/A |
| EM095B | 2029600 | 2024144 | 2288 | 3168 | 0.16 | N/A |
| EM096 | 562860 | 557354 | 1789 | 3717 | 0.66 | N/A |
| EM098 | 446412 | 440438 | 1895 | 4079 | 0.91 | N/A |
| G3679-1 | 1355360 | 1349519 | 1897 | 3944 | 0.29 | N/A |
| G3680-1 | 149784 | 149675 | 13 | 96 | 0.06 | N/A |
| G3682-1 | 3504931 | 3502225 | 615 | 2091 | 0.06 | N/A |
| G3683-1 | 1207038 | 1202736 | 1261 | 3041 | 0.25 | N/A |
| G3686-1 | 4507006 | 4505341 | 559 | 1106 | 0.02 | N/A |
| G3687-1 | 2186658 | 2176377 | 4371 | 5910 | 0.27 | N/A |
| G3729 | 2759764 | 2757795 | 642 | 1327 | 0.05 | N/A |
| G3734-1 | 838143 | 822305 | 7469 | 8369 | 1.00 | N/A |
| G3750-1 | 837699 | 824731 | 5673 | 7295 | 0.87 | N/A |
| G3750-2 | 400166 | 389304 | 3970 | 6892 | 1.72 | N/A |
| G3750-3 | 341177 | 330270 | 4533 | 6374 | 1.87 | N/A |
| G3758 | 1844108 | 1841345 | 1261 | 1502 | 0.08 | N/A |
| G3771 | 341011 | 337765 | 968 | 2278 | 0.67 | N/A |
| G3782 | 1200724 | 1197578 | 599 | 2547 | 0.21 | N/A |
| G3786 | 358637 | 348679 | 2455 | 7503 | 2.09 | N/A |
| G3788 | 1334836 | 1320139 | 5732 | 8965 | 0.67 | N/A |
| G3841 | 971191 | 957371 | 6507 | 7313 | 0.75 | N/A |

Given that we searched only for bacterial 16S sequences it is important to consider that the number of actual sequences from bacteria could be considerably higher since sequences from other bacterial genes would also be expressed in the samples. Figure 1 shows the percentage of the top 30 most abundant bacteria detected in the 99 samples using BBH taxonomic assignment paradigm. There are different bacterial diversity profiles both among samples and patients, although in some cases intra-patient commonalities are recognized. The top 10 most abundant taxa in all the samples (ranked using the average percentage of BBH assigned reads in all the samples) were *uncultured Streptococcus sp., uncultured gamma proteobacterium, uncultured Pseudomonas sp., uncultured actinobacterium, uncultured cyanobacterium, uncultured Firmicutes bacterium, uncultured Staphylococcus sp., uncultured alpha proteobacterium, uncultured beta proteobacterium* and *uncultured Bacilli bacterium.*

**Figure 1:**
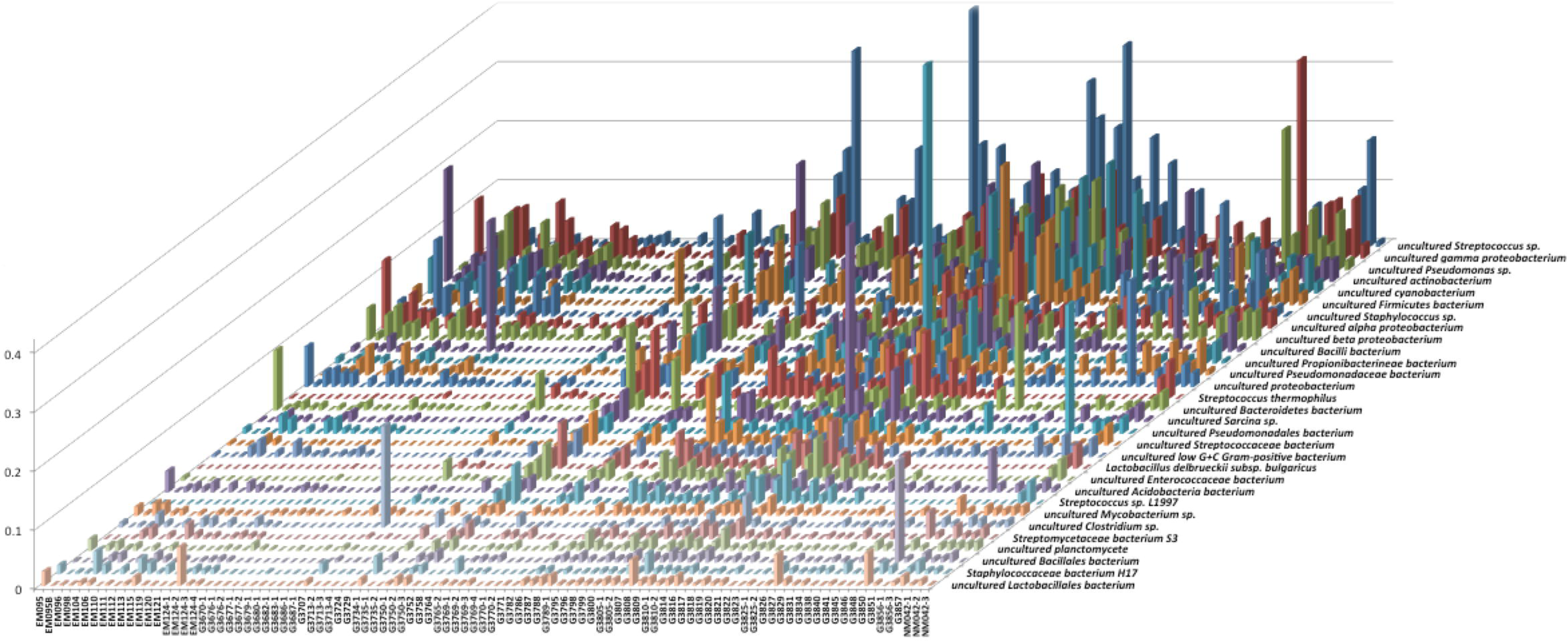
Percentage of the top 30 most abundant bacterial detected in 99 RNA-Seq samples^4^ from Ebola patient serum using the Best Blast Hit paradigm of taxonomic assignment (the frequencies are not cumulative)

Strikingly all these taxa are uncultured bacteria. Symptoms of EVD often mimic those due to bacterial sepsis (http://www.who.int/csr/disease/ebola/clinical-care-summary.pdf). Negative bacterial blood culture in EBV patients with symptoms of bacteraemia or sepsis could be related with the fact that bacteria migth be unculturable. Some samples showed frequency peaks of taxa which were not abundant in the other samples. *Uncultured Streptococcus, uncultured Staphylococcus, uncultured Actinobacterium, uncultured Clostridium or uncultured Bacilli bacterium* were some of these sample-specific peaks.

Figure 2 shows the percentage of the top 30 most abundant bacteria detected using LCA taxonomic assignment paradigm.

**Figure 2:**
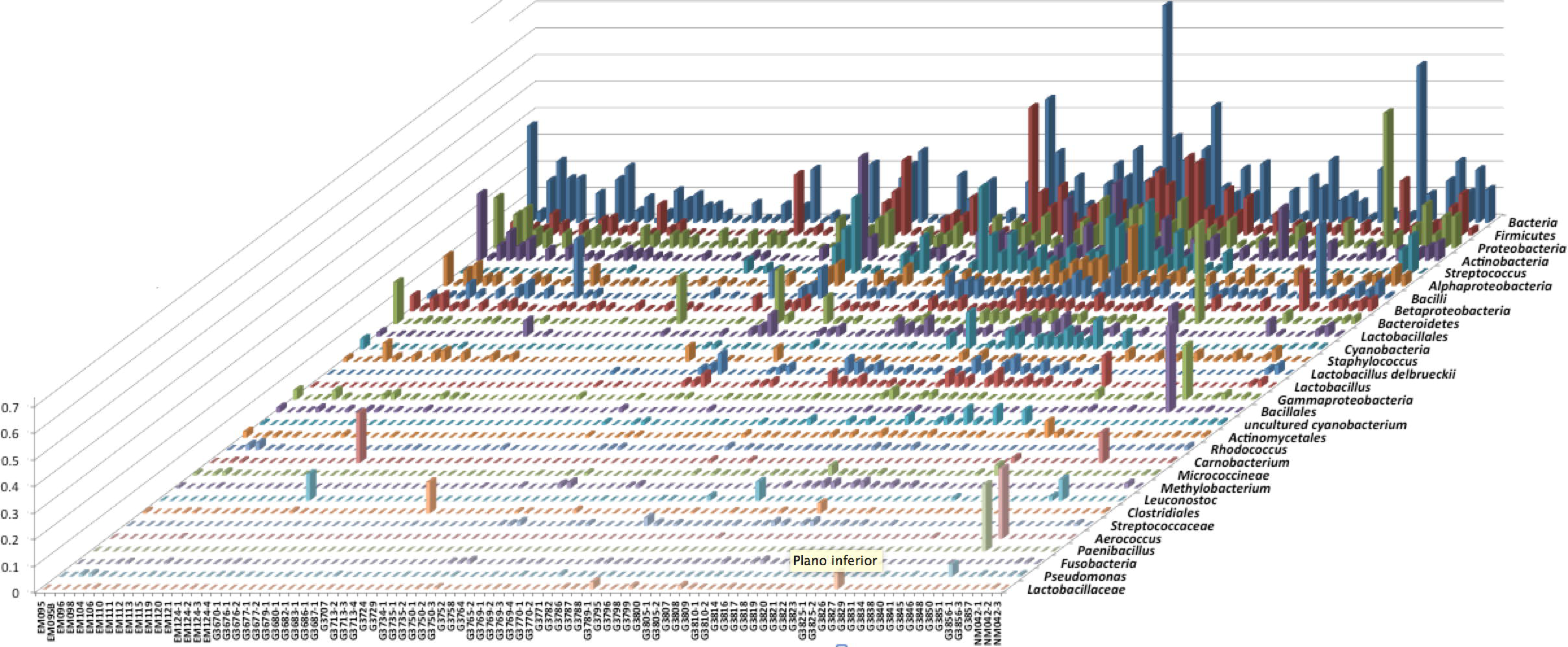
Percentage of the top 30 most abundant bacterial detected in 99 RNA-Seq samples^4^ from Ebola patient serum using the Lowest Common Ancestor paradigm of taxonomic assignment (the frequencies are not cumulative)

Tables and charts with the complete data about the bacteria detected using LCA and BBH paradigms are available as Supplementary material.

## Patient evolution and bacterial load

Tables 2 and 3 and Figures 3 and 4 show EBOV **copies/ml** and **% bacterial reads** in two patients with different outcome. We selected these two patients with 4 available RNA-Seq samples taken at different points of the Ebola disease evolution. Curiously the last point of the discharged patient (G3769-4) shows a very high level of viral EBOV load and very low bacterial load. On the contrary, in the last point of the patient with fatal outcome the bacterial load was higher than the viral EBOV load.

**Figure 3:**
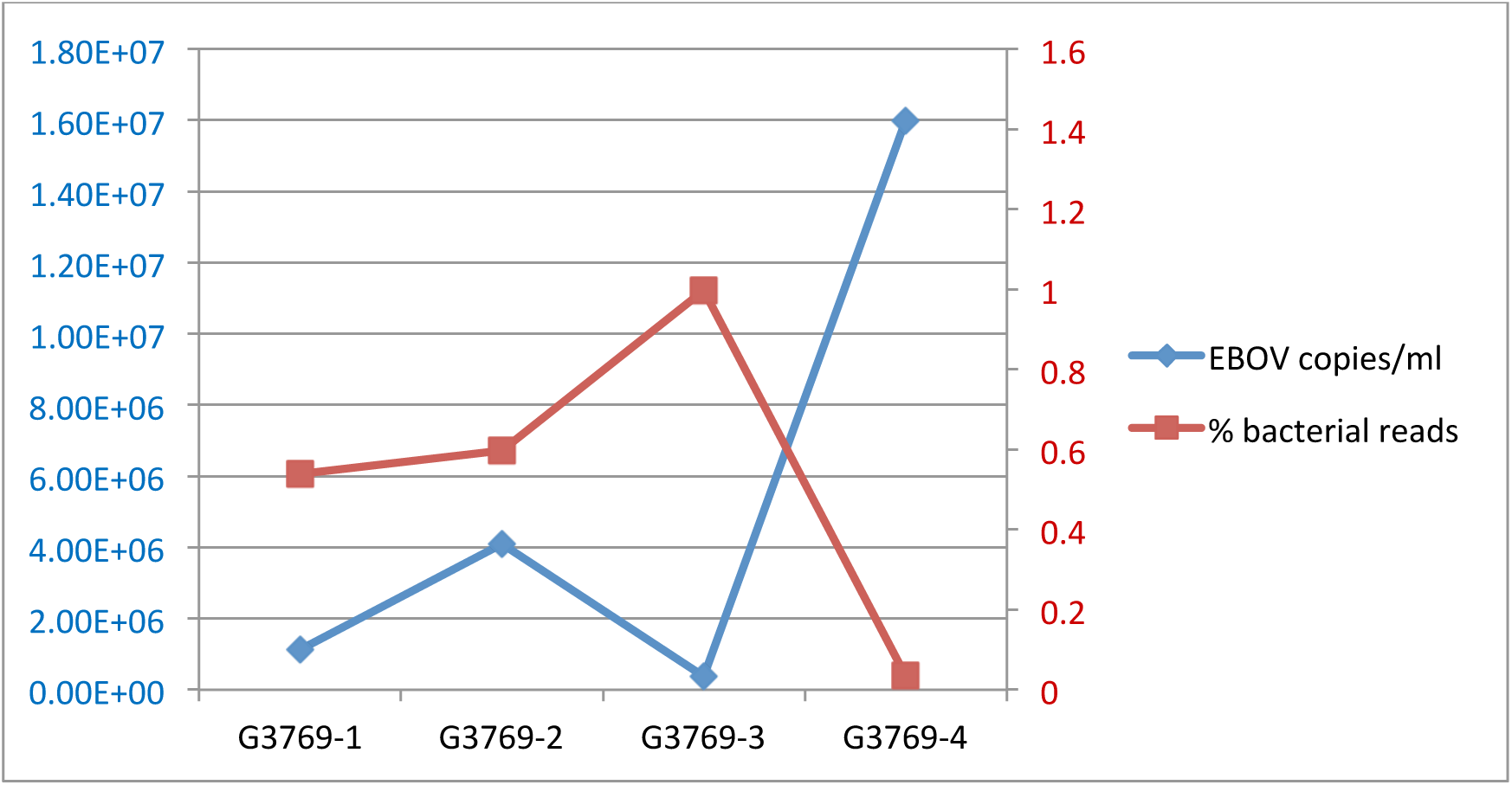
EBOV and bacterial load in 4 evolution points of the Ebola disease in a discharged patient

**Figure 4:**
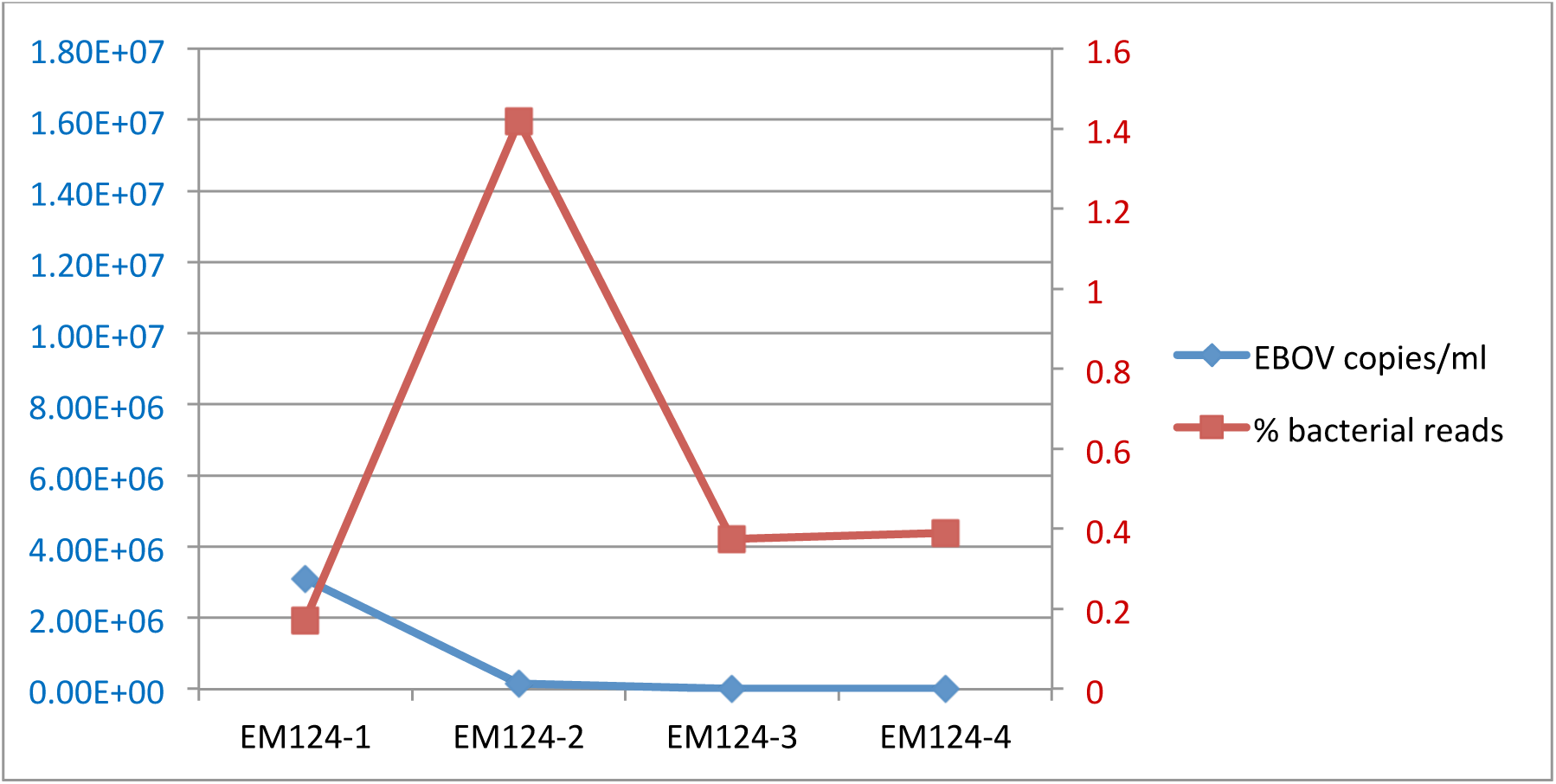
EBOV and bacterial load in 4 evolution points of the Ebola disease in a died patient

**Table 2:** EBOV and bacterial load in 4 evolution points of the Ebola disease in a discharged patient

| Evolution Points | EBOV copies/ml | % bacterial reads | Outcome |
| --- | --- | --- | --- |
| G3769-1 | 1.12E+06 | 0.539633236 | Discharged |
| G3769-2 | 4.10E+06 | 0.597289625 | Discharged |
| G3769-3 | 3.71E+05 | 0.998270488 | Discharged |
| G3769-4 | 1.60E+07 | 0.035288415 | Discharged |

**Table 3:** EBOV and bacterial load in 4 evolution points of the Ebola disease in a patient with fatal outcome

| Evolution points | EBOV copies/ml | % bacterial reads | Outcome |
| --- | --- | --- | --- |
| EM124-1 | 3.11E+06 | 0.171251123 | Died |
| EM124-2 | 1.45E+05 | 1.41980041 | Died |
| EM124-3 | 2.07E+04 | 0.37457228 | Died |
| EM124-4 | 7.44E+03 | 0.389441022 | Died |

It has been reported ^6^ how a patient course was complicated with Gram negative sepsis when the viral load in blood was declining. After treatment with intensive fluid resuscitation, broad-spectrum antimicrobial therapy, and ventilatory support, the patient survived. This finding is coherent with the role that we propose for bacteraemia in EVD.

## New perspective in Ebola disease clinical intervention

This work could open a new perspective in both the pathogenesis and the treatment of EBOV infections. These findings suggest that sepsis therapies, specific antibiotics and mucosal epithelial barrier protectors could be considered. It is possible that, as in other immunocompromised patients, the crucial interventions would be targeting those bacteria. This is especially important in the case of EVD because there is not a specific treatment for Ebola virus.

## Acknowledgments

This work was funded in part by Cardiobiome project ITC-20151148.

